# Mechanical Stretch Counteracts Hypomagnetic Field-Induced Changes in YAP Nuclear Localization in C2C12 Myoblasts

**DOI:** 10.64898/2026.10.13.758941

**Authors:** Joyce Z. Yang, Sachin Kaluarachchi, Kun Man, Gnana Kushal Raju Adidam, Kaavya Akula, Antonio Woollard, Kai Wang

**Author notes:** J.Z.Y. and S.K. contributed equally to this work. Corresponding authors Kai Wang, PhD, 149, 13^th^ St, Room 5.304, Charlestown, MA 02129.

## Abstract

The geomagnetic field (GMF) is a persistent environmental factor that influences biological processes. Reduced geomagnetic field exposure can alter cellular processes, but its effects on cellular mechanotransduction and responsiveness to mechanical stimulation remain poorly understood. Here, we developed an *in vitro* platform combining a programmable uniaxial stretching device with a mu-metal shielding chamber to independently manipulate mechanical stretch and magnetic field exposure. Using C2C12 myoblasts, we evaluated culture conditions and cyclic stretching parameters to identify a regimen that promoted nuclear localization of the mechanosensitive transcriptional coactivator yes-associated protein (YAP). We then compared cell and nuclear morphology and YAP nucleocytoplasmic localization under geomagnetic field (GMF) and hypomagnetic field (HMF; approximately 5 nT) conditions, with or without cyclic stretch. Under static conditions, HMF exposure increased cell and nuclear areas, promoted nuclear elongation, reduced nuclear circularity, and decreased the nuclear YAP level. Cyclic stretch induced nuclear alignment perpendicular to the stretching direction under both magnetic field conditions, without significantly altering cell or nuclear area or nuclear elongation. Importantly, cyclic stretch increased nuclear YAP localization under both GMF and HMF conditions, restoring it in HMF-exposed cells to levels comparable to those in stretched cells under GMF. These findings indicate that C2C12 myoblasts remain responsive to mechanical stimulation under HMF conditions and that cyclic stretch can counteract the HMF-associated reduction in nuclear YAP localization. Our platform provides a foundation for investigating how magnetic and mechanical cues jointly regulate myoblast behavior.

## 1. INTRODUCTION

The geomagnetic field (GMF) is a fundamental component of the Earth’s environment and plays a critical role in supporting life. Beyond its well-established role in animal navigation and magnetoreception,^1^ growing evidence suggests that magnetic fields also affect a wide range of biological processes at the cellular and molecular levels, including DNA and enzymatic activity, cell morphology, differentiation, and organelle functions.^2^ However, magnetic field strength decreases substantially beyond Earth, from approximately 25–65 μT at the surface to 15–50 μT in the low-Earth orbit, and further to hypomagnetic field (HMF; <5 μT) levels on the Moon (300 nT–5 μT), Mars (10–300 nT), and in interplanetary space (0.1–10 nT).^3^ Concerns about the potential health effects of reduced magnetic field exposure date back to the 1960s, particularly in the context of space exploration.^4, 5^ Subsequent studies have reported alterations in brain function, circadian rhythms, and cellular processes such as mitochondrial function under HMF conditions.^3, 6^ Despite these findings, the mechanisms by which reduced magnetic field exposure influences cellular behavior remain poorly understood. Investigating these effects is therefore important not only for understanding the biological challenges of deep-space exploration but also for gaining broader insights into how physical environmental cues regulate cellular function.

Skeletal muscle is particularly relevant in the context of space exploration because its structure and function are highly dependent on mechanical loading. Mechanical stimulation is essential for maintaining muscle structure and function, whereas prolonged mechanical unloading contributes to muscle atrophy and functional decline.^7–9^ At the cellular level, these adaptations are mediated by mechanotransduction, through which cells convert physical forces into biochemical signals that regulate muscle development, maintenance, and regeneration.^10, 11^ Mechanical stretching provides a controlled approach to investigate muscle cell responses to mechanical loading by applying tensile strain to cells. In myoblasts, mechanical stretching has been shown to modulate mechanotransduction signaling and influence cell proliferation and differentiation, with responses depending on culture conditions and stretching regimens.^12, 13^ Importantly, the cellular machinery involved in mechanotransduction, particularly the actin cytoskeleton, may also be sensitive to changes in magnetic field exposure. For example, shielding the GMF alters actin assembly and inhibits motility in human neuroblastoma cells,^6^ while static magnetic field exposure affects proliferation and intracellular protein localization in C2C12 myoblasts.^14^ These findings raise the possibility that changes in magnetic field exposure may influence how skeletal muscle cells sense and respond to mechanical forces. However, whether HMF exposure alters mechanotransduction in skeletal muscle cells remains largely unknown. Furthermore, it is unclear whether skeletal muscle cells maintain their ability to respond to mechanical stimulation under HMF conditions or whether mechanical stretching can counteract HMF-associated alterations in mechanosensitive signaling. Addressing these questions could reveal previously unrecognized interactions between magnetic and mechanical cues in regulating skeletal muscle cell behavior.

In this study, to investigate how HMF affects skeletal muscle cell mechanotransduction in response to mechanical stimulation, we developed an *in vitro* experimental platform that integrates a programmable uniaxial stretching device with a mu-metal shielding chamber, enabling independent control of magnetic field exposure and mechanical stretch. Using C2C12 myoblasts, we first examined cell culture conditions and cyclic stretching regimens to identify conditions that promote nuclear localization of yes-associated protein (YAP), a mechanosensitive transcriptional coactivator that plays a central role in translating mechanical cues into cellular responses. Then, we investigated C2C12 cells under four different conditions: static and GMF, static and HMF, stretch and GMF, and stretch and HMF. Delineating the effects of HMF and mechanical stretching on C2C12 cells provides insight into the role of magnetic field strength in skeletal muscle mechanobiology and approaches to mitigating muscle dysfunction during future deep-space exploration.

## 2. RESULTS AND DISCUSSION

### 2.1. Design and fabrication of experimental apparatus

To investigate the effects of cyclic mechanical stretch and magnetic field on C2C12 cells, we designed and fabricated a motor-driven mechanical stretching device to control cyclic strain and used a mu-metal shielding chamber to create a hypomagnetic field environment.

The mechanical stretching device consists of a stretching unit, a cell culture chamber, and a motor actuator installed on a cell culture base, as shown in **Figure 1**. The stretching unit and cell culture chamber were designed using SolidWorks and 3D-printed using stainless steel. The static and upper clamps and L-shape arm were 3D-printed using resin to facilitate stretching with reduced mass. A flexible polydimethylsiloxane (PDMS) membrane was mounted on the stretching unit, with one side fixed and the other attached to the movable end of the unit that moved along a rail. The movable end was connected to a motor through an L-shaped arm. A rack-and-pinion gear system was utilized to convert the rotational motion of the motor to linear displacement, thereby generating cyclic uniaxial stretch on the membrane. The stretching displacement (strain magnitude) and frequency were precisely controlled by a programmable microcontroller board. The stretching unit was placed inside the cell culture chamber with a glass window at the bottom to facilitate microscopic observation and imaging. To avoid contamination, the entire stretching device was enclosed within a 3D-printed resin container (**Figure 2A**).

**Figure 1.**
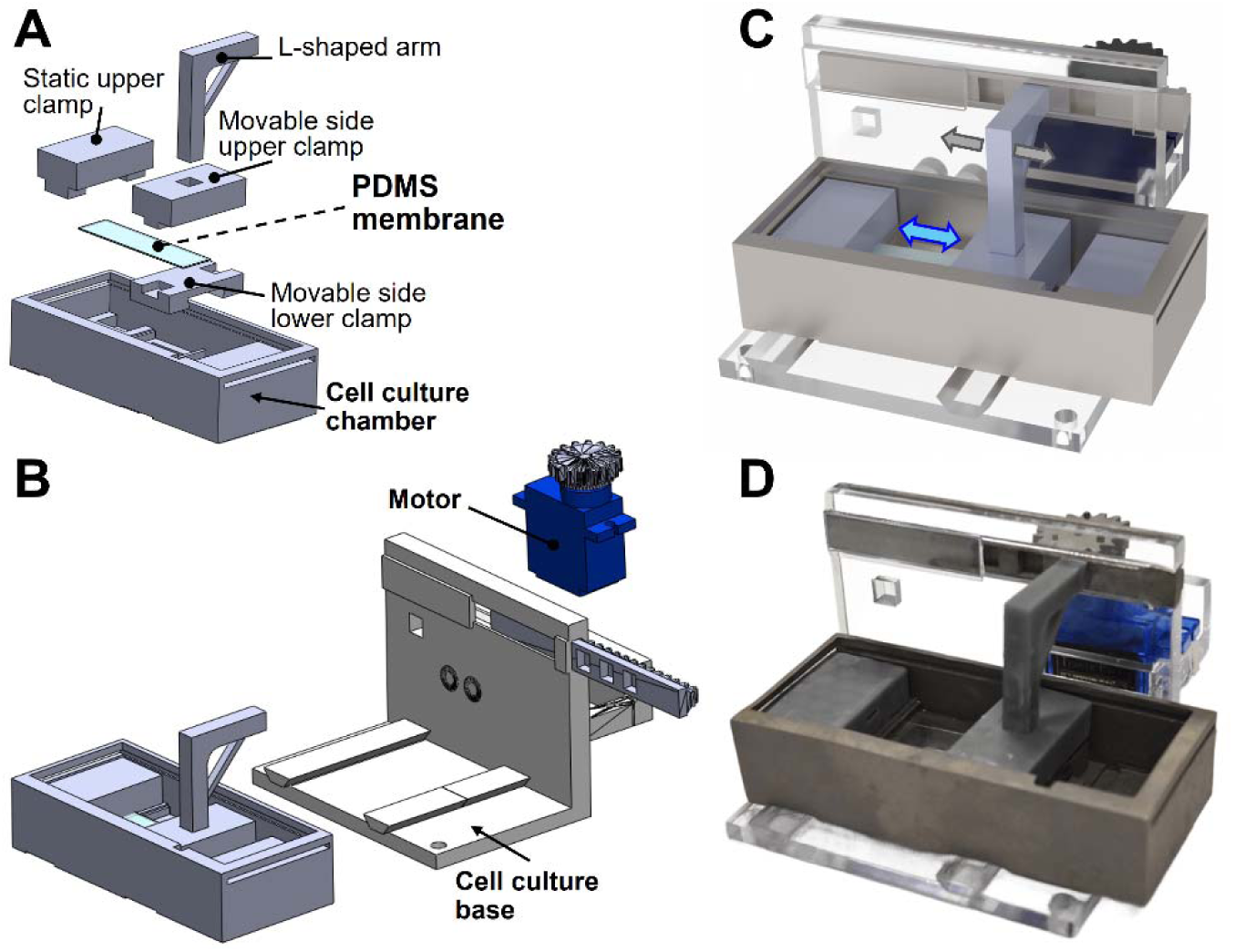
Motor-driven uniaxial stretching device. (**A**) Exploded view of the stretching platform. (**B**) Illustration of an assembled stretching platform with a motor actuator installed on a cell culture base. (**C**) Illustration and (**D**) photo of a uniaxial stretching device.

**Figure 2.**
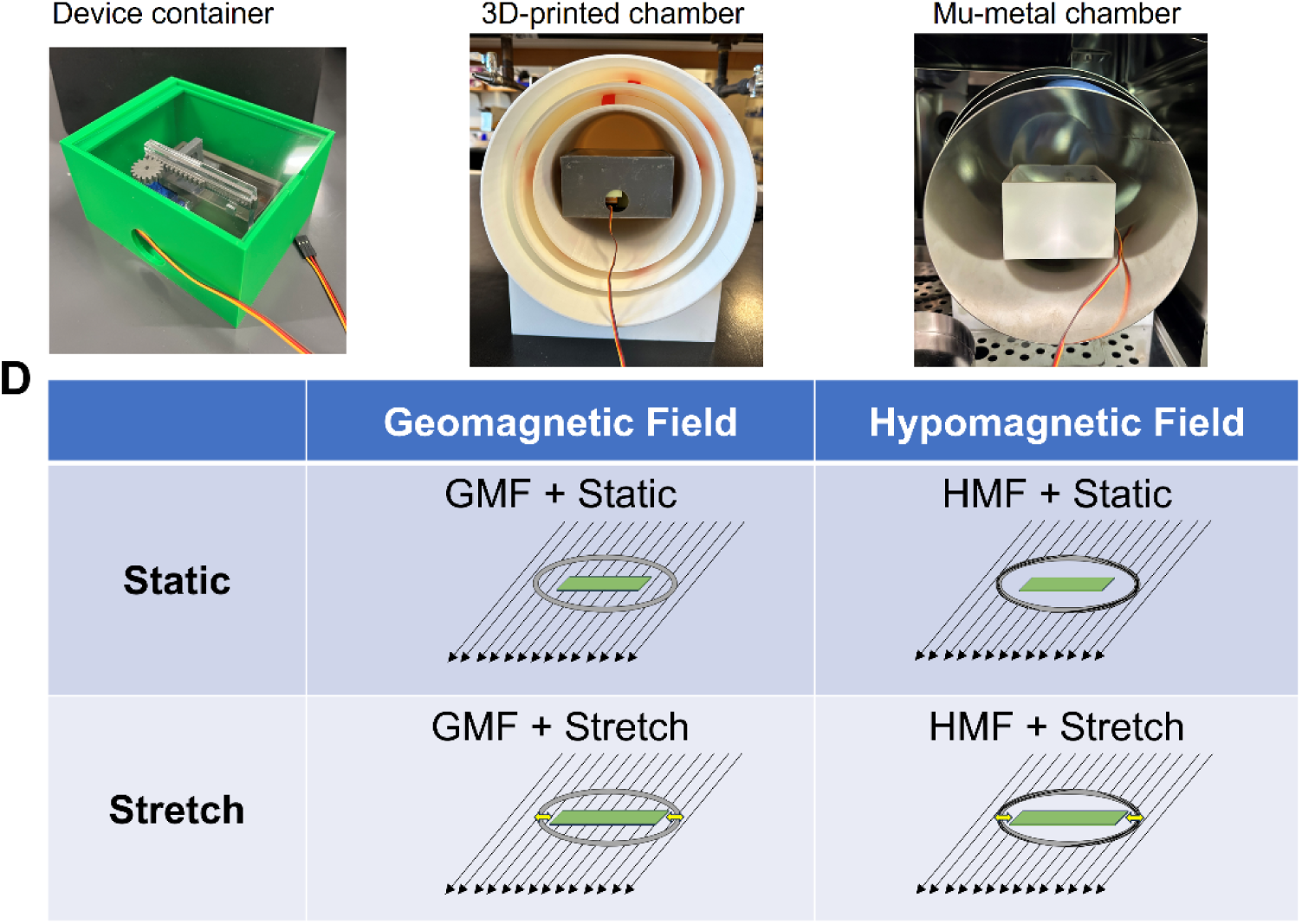
Experimental apparatus and design. (**A**) Photograph of a stretching device placed inside a 3D-printed plastic container. (**B**) 3D-printed plastic chamber with the same dimensions as the mu-metal chamber. (**C**) mu-metal chamber housing a stretching device. (**D**) Schematic of experimental design.

C2C12 myoblasts were cultured on a PDMS membrane within the stretching device and either maintained under static conditions (Static) or subjected to cyclic stretching (Stretch) under different magnetic field conditions. A mu-metal shielding chamber was used to attenuate the ambient geomagnetic field by redirecting magnetic flux through its high-permeability walls, creating a hypomagnetic field (HMF) of approximately 5 nT.^15^ To match the culture condition, a 3D-printed plastic chamber with identical dimensions was used in the GMF control group. Together, the stretching device and magnetic shielding chambers enabled independent manipulation of magnetic and mechanical cues to investigate their individual and combined effects on mechanotransduction in C2C12 myoblasts (**Figure 2**).

### 2.2. Optimizing cell culture conditions and cyclic stretching regimen

Previous studies have shown that myoblast responses to mechanical stimulation are highly sensitive to culture conditions, stretching modes (uniaxial, equibiaxial, or radial), strain magnitude, and stimulation regimens.^16^ Therefore, establishing appropriate cell culture conditions and stretching parameters is essential for reliably evaluating the effects of magnetic field exposure on cellular mechanotransduction. To assess cellular responses to mechanical stimulation, we focused on evaluating yes-associated protein (YAP). YAP acts as a mechanosensitive transcriptional coactivator regulated by the Hippo signaling pathway and plays an important role in myogenesis, skeletal muscle homeostasis, and muscle disorders. Exercise and mechanical overload have been shown to modulate YAP expression and phosphorylation in mouse skeletal muscle,^17, 18^ while altered YAP activity has been associated with both muscle atrophy and hypertrophy.^19–21^ In cultured skeletal muscle cells, elevated YAP expression and activity promote cell proliferation, whereas YAP inactivation favors myogenic differentiation.^22, 23^ Given its sensitivity to mechanical cues, we used YAP nuclear localization as a readout of mechanotransduction to evaluate cell culture conditions and optimize cyclic stretching parameters.

First, we examined the concentration of type I collagen coating on the PDMS membrane surface. Increasing collagen concentration from 25 to 50 to 100 µg/mL did not significantly change cell morphology or YAP subcellular localization (**Figure 3A**-**C**). On the PDMS membrane coated with 50 µg/mL collagen, a higher cell seeding density (20,000 cells/cm^2^) led to more transitional YAP expression (nuclear and cytoplasmic YAP expression is not distinct) (**Figure 3D** vs. **3B**). Because increased cell–cell interactions could interfere with the effects of mechanical stretch,^24^ making it challenging to analyze the effects of mechanical stretch on cell behavior. We therefore selected 5,000 cells/cm^2^ seeding density on the PDMS membrane coated with 50 µg/mL collagen for downstream studies.

**Figure 3.**
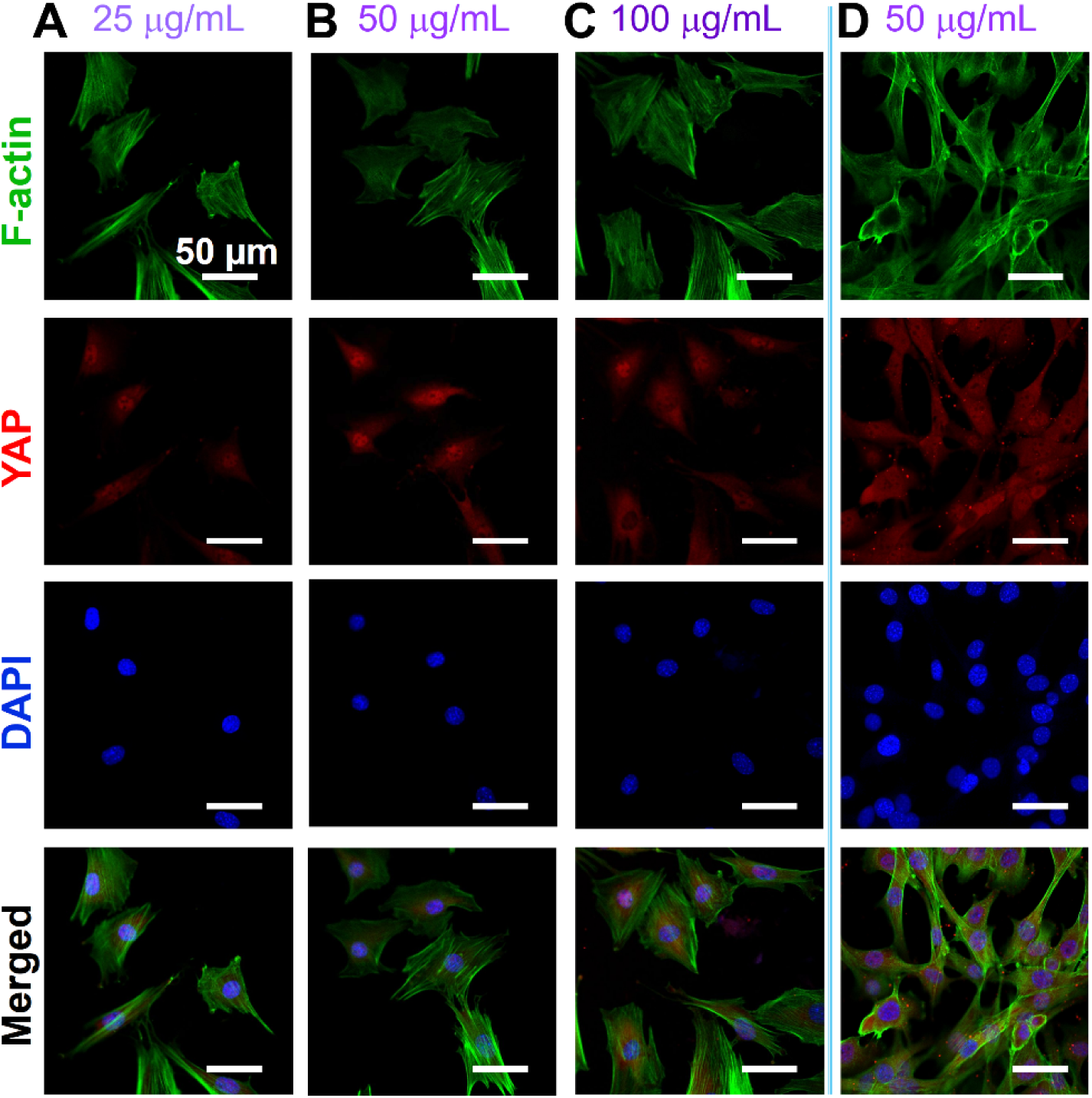
Effects of collagen coating concentration and cell seeding density on YAP expression in C2C12 cells under static conditions. Representative immunofluorescence images of C2C12 cells cultured on PDMS membranes coated with (**A**) 25 µg/mL, (**B**, **D**) 50 µg/mL, (**C**) 100 µg/mL collagen type I. Cell seeding density was 5,000 cells/cm² in (**A**–**C**) and 20,000 cells/cm² in (**D**). Cells were stained for F-actin (green), YAP (red), and nuclei with DAPI (blue). Scale bars: 50 µm.

Cyclic mechanical stretch at frequencies of 0.5–2 Hz and strain magnitudes of 5–20% has been widely applied to investigate the responses of C2C12 myoblasts to mechanical stimulation.^25–27^ Building on these established studies, we evaluated cyclic stretch at 5–10% linear strain and 0.5–1 Hz to determine how stretching parameters influence YAP subcellular localization in C2C12 cells. Other important parameters we assessed were the stretch duration and the rest interval between stretches. We first applied continuous cyclic stretch at 5% strain and 1 Hz for 2 h, under which most cells exhibited predominantly cytoplasmic or mixed nuclear– cytoplasmic YAP localization (**Figure 4A**). Extending the stimulation regimen to two 2-h stretching periods separated by a 30-min rest interval further reduced nuclear YAP localization (**Figure 4B**), suggesting that prolonged stretching may attenuate the YAP nuclear response. We therefore shortened each stretching period to 1 h while maintaining 5% strain, 1 Hz frequency, and a 30-min rest interval. This modification resulted in more pronounced nuclear YAP localization (**Figure 4C**). Further reducing the rest interval from 30 to 10 min decreased cytoplasmic YAP staining (**Figure 4D**). In contrast, lowering the stretching frequency from 1 to 0.5 Hz reduced the proportion of cells exhibiting nuclear YAP localization (**Figure 4E**). Collectively, these observations indicate that YAP subcellular localization is sensitive to stretching duration, rest intervals, and frequency, highlighting the importance of optimizing mechanical stimulation regimens to elicit a robust mechanotransduction response.

**Figure 4.**
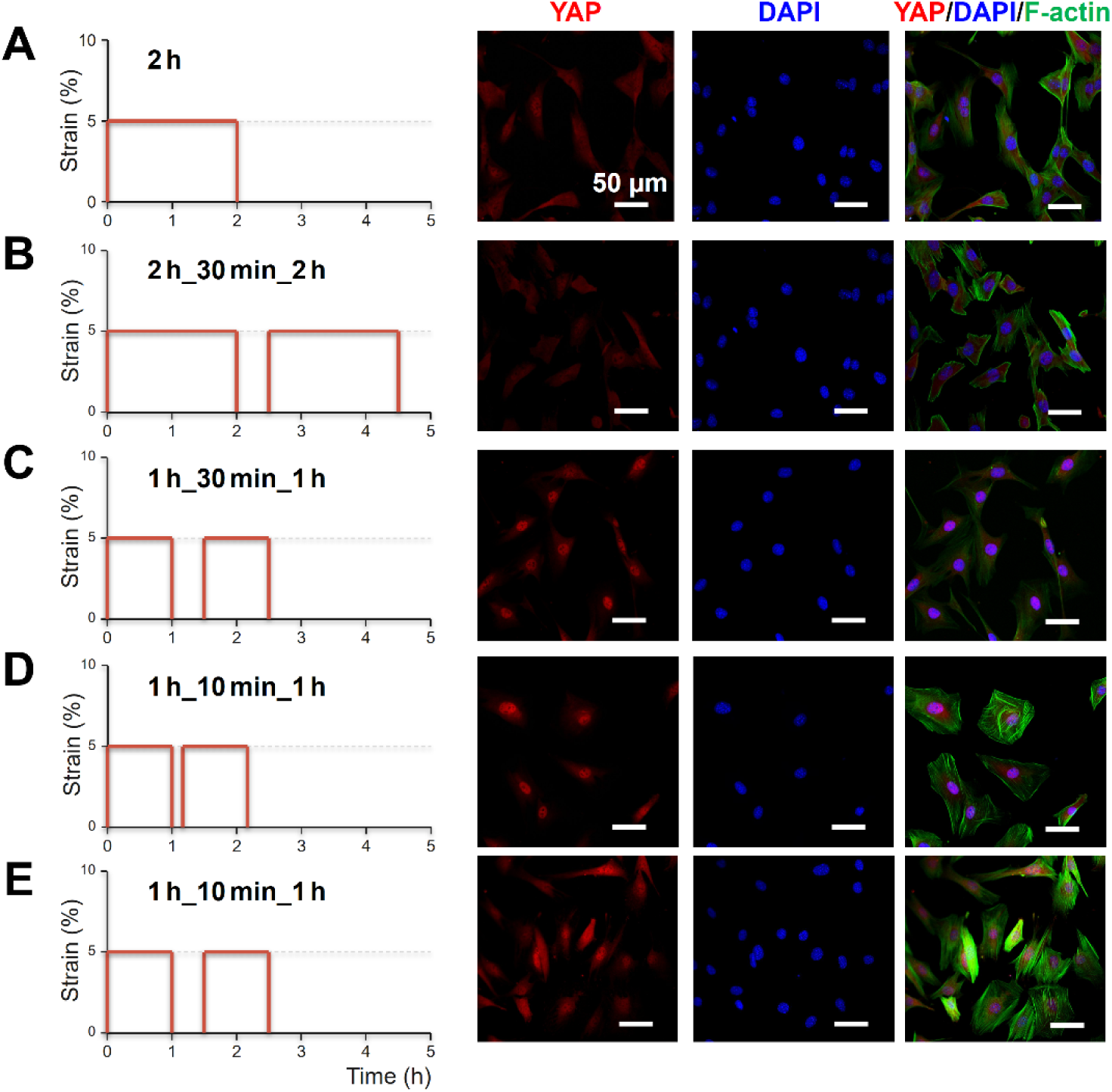
Effects of cyclic stretching regimens on YAP expression in C2C12 cells. C2C12 cells cultured on PDMS membranes coated with 50 µg/mL collagen type I were subjected to cyclic stretch at 5% linear strain and 1 Hz under different regimens: (**A**) 2 h of continuous stretch; (**B**) 2 h of stretch followed by 0.5 h of rest and another 2 h of stretch; (**C**) 1 h of stretch followed by 30 min of rest and another 1 h of stretch; and (**D**) 1 h of stretch followed by 10 min of rest and another 1 h of stretch. (**E**) Cells were subjected to 1 h of stretch followed by 10 min of rest and another 1 h of stretch at 5% strain and 0.5 Hz. Representative immunofluorescence images show YAP (red), DAPI (blue), and F-actin (green). Scale bars: 50 µm.

Having established that cyclic stretch at 5% strain at 1 Hz for two 1 h periods, separated by a 10 min rest interval (abbreviated as 1 h_10 min_1 h stretch), significantly increased nuclear YAP localization compared with static control, we next examined whether increasing strain magnitude could further enhance nuclear YAP localization. To assess the effects of strain magnitude under identical culture conditions, we generated a strain gradient of 2.5–35% on a tapered PDMS membrane as previously reported.^28^ As shown in **Figure 5**, cyclic stretch at 2.5–5.5% strain significantly increased the YAP nuclear-to-cytoplasmic (nuc/cyto) ratio compared with static conditions. However, further increases in strain magnitude reduced the YAP nuc/cyto ratio.

**Figure 5.**
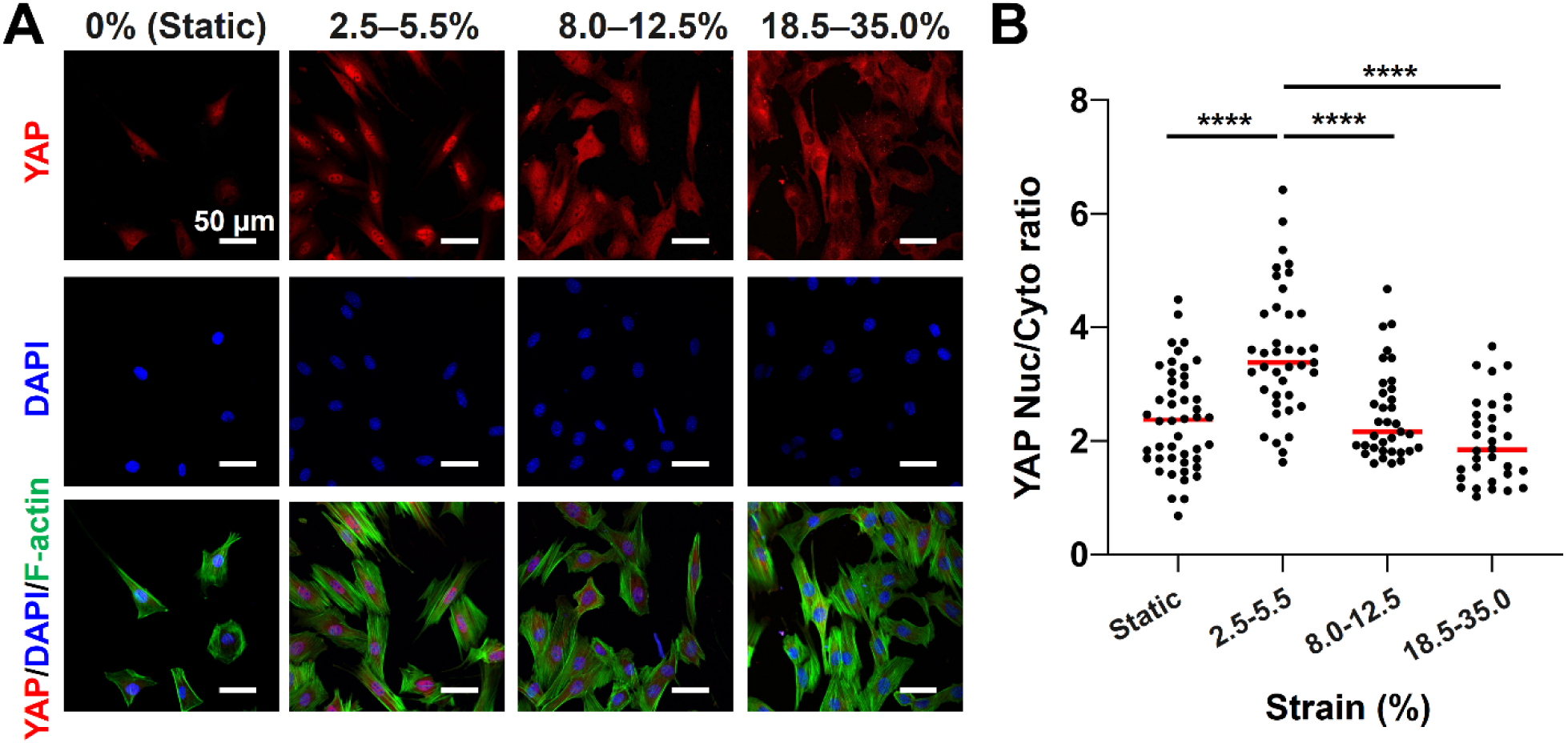
Effects of strain magnitude on YAP expression in C2C12 cells. C2C12 cells cultured on tapered PDMS membranes coated with 50 μg/mL collagen I were subjected to cyclic stretch at 1 Hz for 2 h with 10 min interval. (**A**) Representative immunofluorescence images of C2C12 cells stained for F-actin (green), YAP (red), and nuclei with DAPI (blue). Scale bars: 50 μm. (**B**) Quantitative analysis of the YAP nuc/cyto ratio in C2C12 cells subjected to different strain ranges. n = 31–56. *: *p* < 0.05, **: *p* < 0.01, ***: *p* < 0.005, ****: *p* < 0.001.

Notably, at 18.5–35.0% strain, the YAP nuc/cyto ratio was lower than under static conditions, and a substantial proportion of C2C12 cells exhibited cytoplasmic YAP localization, compared with more transitional localization at 8–12.5% strain and predominantly nuclear localization at 2.5–5.5% strain (**Figure 5A**). After identifying the stretching regimen, we then examined how total stretching duration affected YAP localization. Using the selected regimen, we extended the total stretching duration from 2 h to 4 and 6 h. As shown in **Figure 6**, longer stretching durations increased nuclear YAP localization. Taken together, we selected cyclic stretch at 5% strain and 1 Hz using the 1 h_10 min_1 h regimen, with a total duration of 4 h, for subsequent HMF studies.

**Figure 6.**
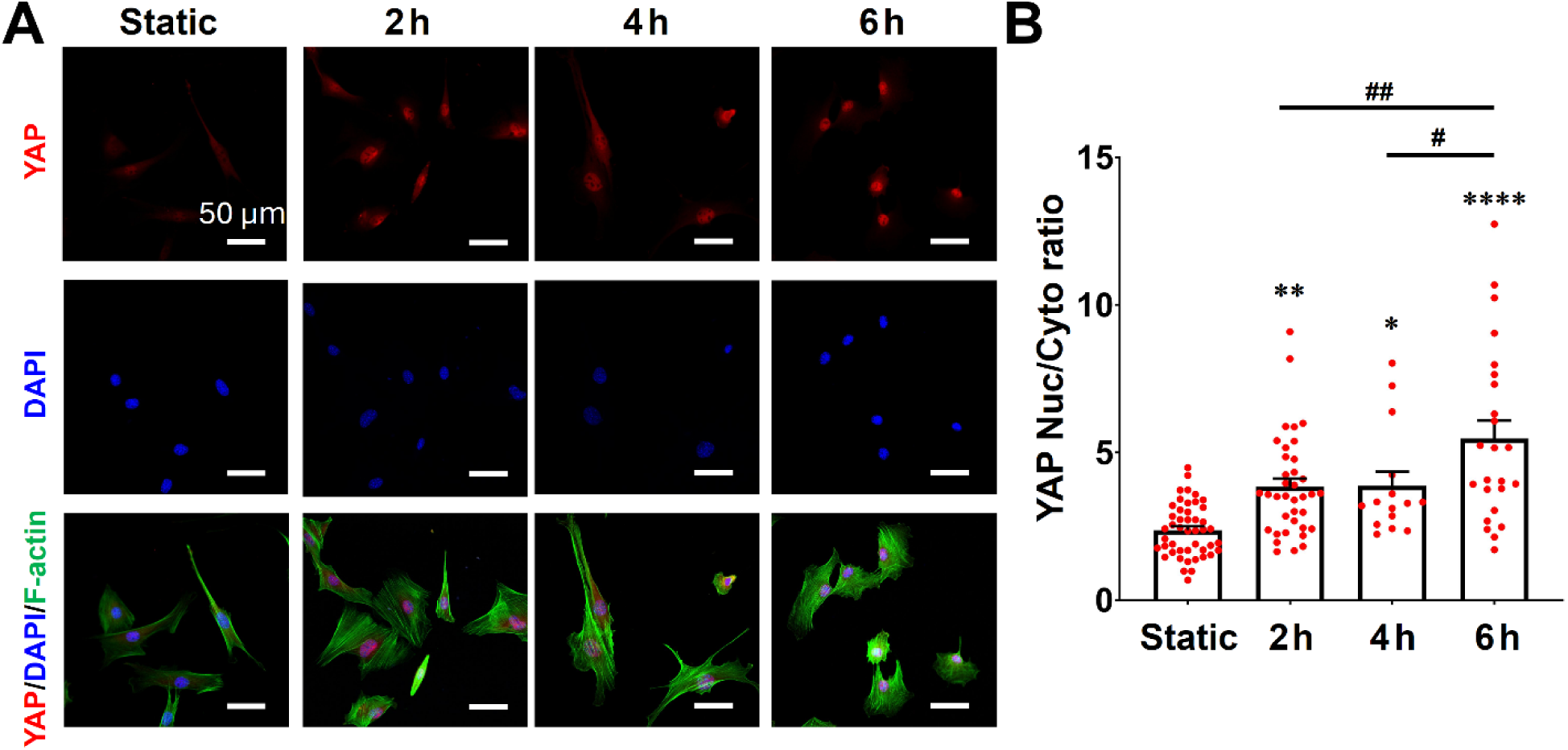
Effects of stretch duration on nuclear YAP expression in C2C12 cells. Cells cultured on PDMS membranes coated with 50 μg/mL collagen I were subjected to cyclic stretch at 5% linear strain and 1 Hz for different duration, with a 10 min rest interval between consecutive 1 h stretching periods. (**A**) Representative immunofluorescence images of C2C12 cells stained for YAP (red), F-actin (green), and nuclei with DAPI (blue). Scale bars: 50 μm. (**B**) Quantitative analysis of the YAP nuc/cyto ratio in C2C12 cells subjected to different stretch durations compared with static controls (2 h without stretch). n = 32–46. *: *p* < 0.05, **: *p* < 0.01, ***: *p* < 0.005, ****: *p* < 0.001.

### 2.3. Delineating the effects of magnetic fields and cyclic stretch on C2C12 cells

Using the optimized culture conditions and stretching parameters, we next investigated how magnetic field exposure and cyclic mechanical stretch influence C2C12 cell and nuclear morphology. Under static conditions, HMF exposure significantly increased both cell and nuclear areas, whereas cyclic stretch had no significant effect on either parameter under GMF or HMF conditions (**Figure 7B, C**). Interestingly, cyclic stretch increased the proportion of nuclei oriented within 60°–90° relative to the stretching axis, indicating preferential nuclear alignment perpendicular to the stretch direction under both magnetic field conditions (**Figure 7D**). This observation is consistent with previous studies showing that myoblasts subjected to cyclic uniaxial stretch tend to reorient perpendicular to the stretching direction, depending on substrate topography, stiffness, and stretching parameters.^29–31^ Such reorientation has been attributed to cytoskeletal remodeling toward the direction of minimum strain.^32^ Our findings suggest that this stretch-associated alignment response is preserved under HMF conditions. HMF exposure increased nuclear elongation and reduced nuclear circularity under static conditions, while cyclic stretch had no significant effect on these nuclear shape parameters (**Figure 7E,F**). Together, these findings suggest that HMF exposure influences cell size and nuclear morphology, whereas cyclic mechanical stretch predominantly affects nuclear orientation, highlighting distinct cellular responses to magnetic and mechanical cues.

**Figure 7.**
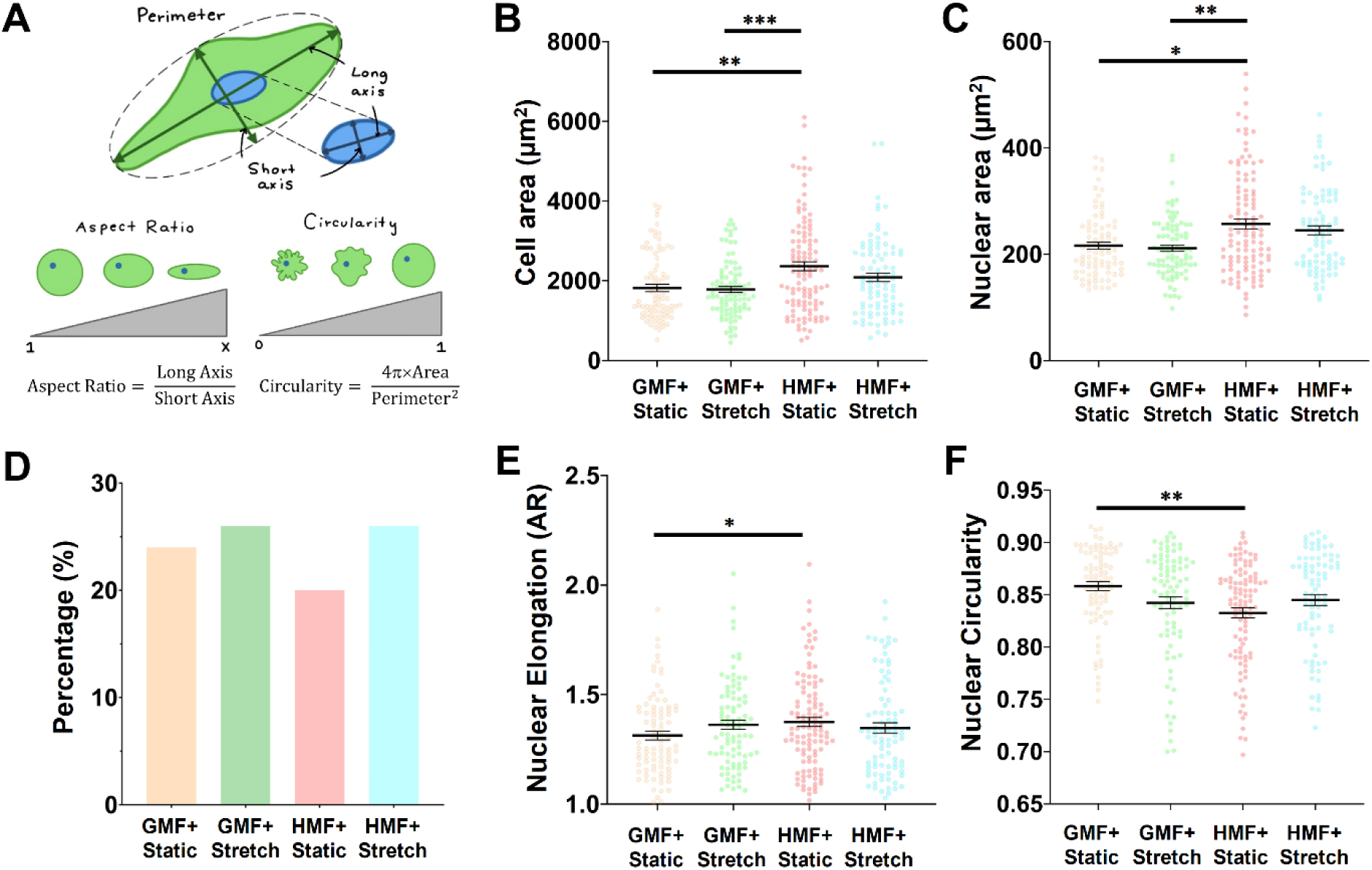
Effects of magnetic field and mechanical stretch on cell and nuclear morphology. (**A**) Definitions of aspect ratio (elongation) and circularity. Quantification of (**B**) cell area, (**C**) nuclear area, (**D**) nuclear alignment, (**E**) nuclear elongation (AR), and (**F**) nuclear circularity under different magnetic field and stretching conditions. In (**D**), percentages indicate the proportion of nuclei oriented within 60°– 90° relative to the stretch direction (toward the perpendicular orientation). *n* = 90–115 cells per condition. * *p* < 0.05, ** *p* < 0.01, *** *p* < 0.005.

We next investigated whether HMF exposure and cyclic mechanical stretch influence YAP nucleocytoplasmic localization in C2C12 myoblasts. Under static conditions, HMF exposure reduced nuclear YAP localization, whereas cyclic stretch increased nuclear YAP localization under both GMF and HMF conditions, counteracting the HMF-associated reduction and bringing nuclear YAP levels to those comparable to stretched cells under GMF (**Figure 8**). Previous studies have shown that repeated mechanical stretching can promote cell spreading and stress fiber formation, even on soft substrates,^33^ while increased cell spreading may facilitate YAP nuclear translocation by enhancing nuclear pore permeability.^34, 35^ However, despite the stretch-induced increase in nuclear YAP localization, we observed no significant changes in cell or nuclear morphology following cyclic stretch (**Figure 7**). Moreover, HMF exposure increased cell and nuclear areas, as well as nuclear elongation, under static conditions, contrasting with previous reports of HMF-induced reductions in cell size and disruptions in F-actin organization.^6^ Notably, these morphological changes were not consistently associated with YAP nuclear localization, suggesting that changes in overall cell or nuclear shape alone cannot fully explain the observed YAP responses. Given that mechanotransduction depends not only on cell morphology but also on the organization and dynamics of the actin cytoskeleton,^36^ alterations in cytoskeletal architecture and associated signaling pathways may contribute to the effects of HMF and cyclic stretch on YAP nucleocytoplasmic localization. Further investigation is needed to elucidate the underlying mechanisms.

**Figure 8.**
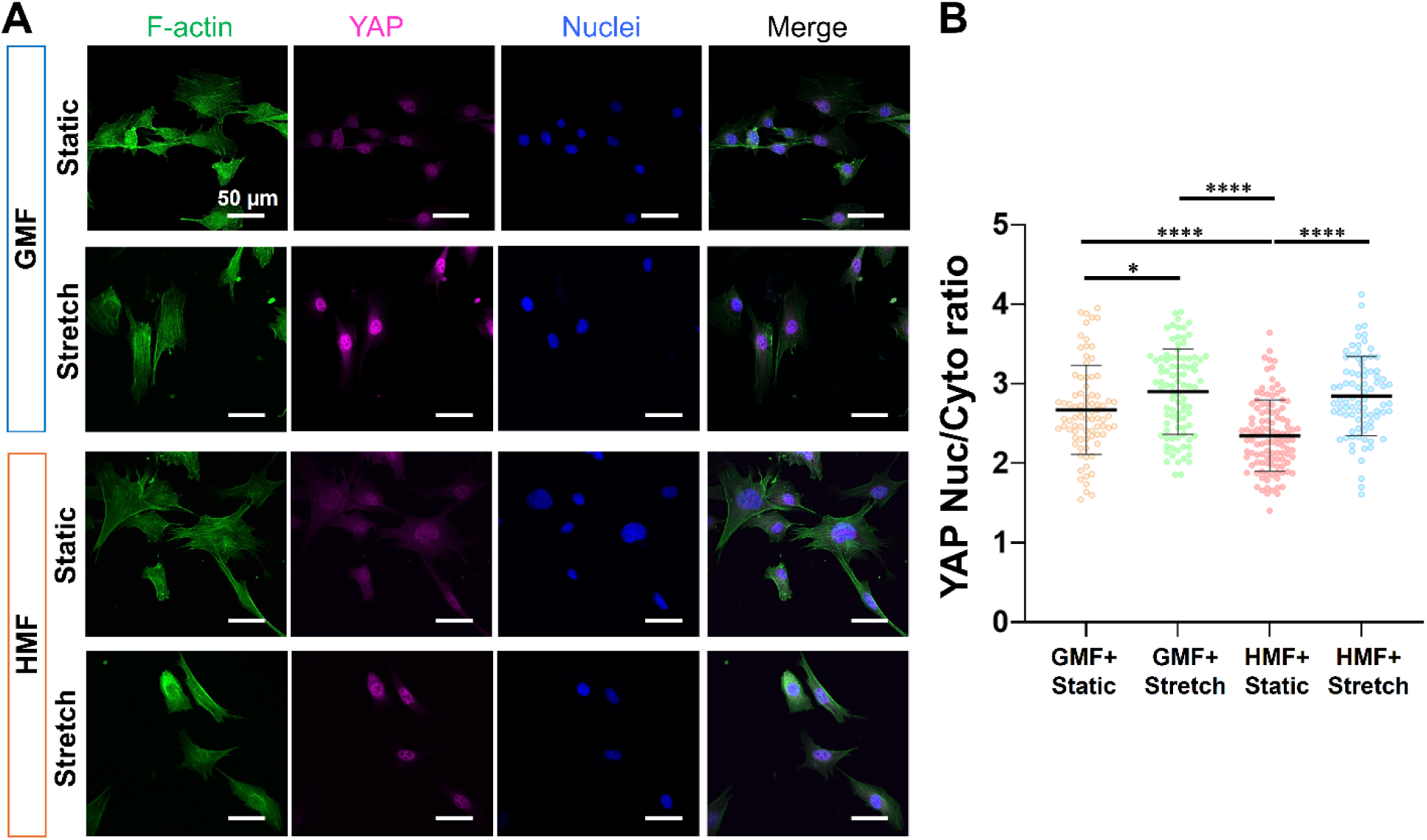
Effects of HMF and mechanical stretch on nuclear YAP expression in C2C12 cells. Cells cultured on PDMS membranes coated with 50 mg/mL collagen I were subjected to cyclic stretch at 5% linear strain and 1 Hz for two 1 h periods separated by a 10 min rest interval. (**A**) Representative immunofluorescence images of C2C12 cells stained for F-actin (green), YAP (purple), and nuclei with DAPI (blue). Scale bars: 50 μm. (**B**) Quantitative analysis of the YAP nuc/cyto ratio of C2C12 cells under different conditions. n = 90–115. *: *p* < 0.05, **: *p* < 0.01, ***: *p* < 0.005, ****: *p* < 0.001.

## 3. CONCLUSION

In this study, we established an experimental platform integrating a programmable uniaxial stretching device with magnetic shielding to investigate the individual and combined effects of HMF exposure and cyclic mechanical stretch on mechanotransduction in C2C12 myoblasts. We identified culture conditions and stretching parameters that promoted nuclear YAP localization. HMF exposure altered cell and nuclear morphology and reduced nuclear YAP localization under static conditions, whereas cyclic stretch counteracted this reduction and promoted nuclear alignment under both GMF and HMF conditions. These findings demonstrate that myoblasts retain responsiveness to mechanical stimulation under HMF conditions and that cyclic stretch can modulate HMF-associated changes in YAP nucleocytoplasmic localization. This platform provides a foundation for elucidating how magnetic and mechanical cues regulate cellular mechanotransduction and for evaluating mechanical stimulation as a potential strategy to mitigate HMF-associated cellular alterations.

## 4. MATERIALS AND METHODS

### 4.1. Fabrication of Mechanical Stretching Device

The stretching platform and cell culture chamber were designed using SolidWorks (Dassault Systèmes SolidWorks Co.) and fabricated by 3D printing using resin and stainless steel, respectively (Unionfab, Shanghai, China). The membrane was stretched using a motor (SG90 9G micro servo motor, SIPYTOPF Co.) connected to an L-shaped arm, controlled by an Arduino board (MEGA R3 Board ATmega 2560, ELEGOO Co.). To minimize contamination, the entire device was enclosed within a 3D-printed resin container.

### 4.2. Preparation of Stretchable Membranes

Two types of flexible polydimethylsiloxane (PDMS) membranes, uniform and tapered, were fabricated using Sylgard 184 (Dow Corning; base-to-crosslinker ratio = 10:1, w/w). To prepare uniform PDMS membranes, the PDMS mixture was cast or spin-coated onto a glass slide using a spin coater (Laurell WS-650MZ-23NPPB), followed by curing at 110 °C for 20 min. The resulting membranes had a thickness of 100–150 µm. To prepare tapered PDMS membranes, a glass coverslip (2.4 cm × 4 cm × 0.15 mm) was placed over a PDMS film spin-coated onto a plastic substrate, with one end of the coverslip contacting the substrate and the other end elevated by a stack of two 0.15-mm-thick glass coverslips. The assembly was then cured at 75 °C for 1 h. The resulting membranes were trimmed into rectangles measuring approximately 40 mm × 8 mm. The membranes were then attached to the stretching devices and treated with oxygen plasma at medium power for 30 s using a plasma cleaner (Model PDC-001, Harrick Plasma, Ithaca, NY, USA), followed by sterilization in ethanol for 30 min. Prior to cell seeding, the PDMS membranes were coated with collagen type I (rat tail; Cat. No. 354236; Corning) at various concentrations (20, 50, and 100 µg/mL) overnight in a cell culture incubator, followed by three washes with phosphate-buffered saline (PBS; Cat. No. 10010023; Thermo Fisher Scientific).

### 4.3. Cell Culture

Mouse C2C12 myoblasts (Cat. No. CRL-1772; ATCC) were cultured in Dulbecco’s modified Eagle medium (DMEM; Cat. No. 30-2002; ATCC) supplemented with 10% fetal bovine serum (FBS; Cat. No. 30-2020; ATCC) and 1% penicillin-streptomycin (Cat. No. 30-2300; ATCC). Cells were maintained at 37 °C in a humidified incubator with 5% CO. Unless otherwise specified, C2C12 cells were seeded onto collagen type I-coated PDMS membranes at a density of 5,000 cells/cm² and cultured overnight before exposure to GMF or HMF conditions, with or without mechanical stretching.

### 4.4. Mechanical Stretching

Two hours before each experiment, the mu-metal shielding chamber and control chamber were placed in a cell culture incubator to equilibrate to the experimental temperature. All components of the stretching devices were disinfected with 70% ethanol and exposed to UV light for 30 min. The Arduino microcontroller was programmed to maintain a constant strain amplitude throughout the experiment. Applied strain was calculated as the change in PDMS membrane length relative to its initial length and verified using caliper measurements. Following assembly, the strain level of each stretching device was checked to ensure consistency across experiments.

Four devices were then placed into their respective enclosures, with one stretching device and one static control housed in the mu-metal chamber and the remaining two devices housed in the control chamber. Static controls were subjected to identical handling and environmental conditions as the stretched groups, except that no mechanical stretching was applied. Cyclic mechanical stretching was performed at the designated strain levels and durations. Immediately following treatment, cells were washed with PBS and fixed with 4% paraformaldehyde (PFA; Cat. No. 047377.9L; Thermo Fisher Scientific).

### 4.5. Hypomagnetic Field Exposure

HMF conditions (∼5 nT) were achieved using a MuMETAL® Zero Gauss Chamber (Magnetic Shield Corp.), which attenuates external static and low-frequency alternating magnetic fields by up to 1,000,000-fold. According to the manufacturer’s specifications, the chamber provides residual magnetic fields below 5 nT after degaussing, with low magnetic noise levels (less than 30 fT/√Hz).

### 4.6. Immunofluorescence Staining

C2C12 cells were fixed with 4% PFA for 20 min at room temperature and washed three times with PBS. Cells were then permeabilized and blocked in PBS containing 1% bovine serum albumin (BSA; Akron Biotech, Boca Raton, FL, USA), 10% goat serum (Cat. No. 16210064; Gibco), and 0.2% Triton X-100 (Cat. No. X100-5ML; Sigma-Aldrich) for 1 h at room temperature. Subsequently, samples were incubated overnight at 4 °C with a primary antibody against YAP (Cat. No. sc-101199; Santa Cruz Biotechnology; 1:100 dilution) and Alexa Fluor 488-conjugated phalloidin (Cat. No. A12379; Invitrogen; 1:100 dilution) in the same blocking solution. After three washes with PBS, cells were incubated with the corresponding secondary antibody diluted in blocking solution for 1 h at room temperature, followed by three additional PBS washes. Samples were then mounted using SlowFade Diamond Antifade Mountant with DAPI (Cat. No. S36968; Invitrogen) for nuclear counterstaining. Fluorescence images were acquired using a Zeiss 710 confocal microscope or a Nikon AXR confocal microscope.

### 4.7. Image Analysis

Cell alignment and nuclear morphology, including circularity and elongation, were quantified using ImageJ (National Institutes of Health, Bethesda, MD, USA). YAP nuclear localization was quantified by calculating the nuclear-to-cytoplasmic (nuc/cyto) fluorescence intensity ratio, following a previously reported method.^37^ Nuclear regions were identified using DAPI staining and the “Analyze Particles” tool, while whole-cell boundaries were delineated based on F-actin staining. The area and integrated fluorescence intensity (IntDen) of YAP were measured separately within the nuclear and whole-cell regions. Mean nuclear YAP fluorescence intensity was calculated by dividing the nuclear IntDen by the nuclear area. Mean cytoplasmic YAP fluorescence intensity was calculated by subtracting the nuclear IntDen and area from the corresponding whole-cell measurements, respectively, and dividing the resulting cytoplasmic IntDen by the cytoplasmic area. The YAP nuc/cyto ratio was then calculated by dividing the mean nuclear fluorescence intensity by the mean cytoplasmic fluorescence intensity.

### 4.8. Statistical Analysis

Statistical analyses were performed using GraphPad Prism (GraphPad Software, San Diego, CA, USA). Data are presented as mean ± standard error of the mean (SEM). Differences among groups were assessed using a one-way analysis of variance (ANOVA), followed by Tukey’s multiple comparisons test. A *p* value < 0.05 was considered statistically significant.

## 5. ACKNOWLEDGEMENT

This work was supported by funding from the Discovery Center for Musculoskeletal Recovery at the Schoen Adams Research Institute, Spaulding Rehabilitation Hospital, and the National Institute of Arthritis and Musculoskeletal and Skin Diseases (R21AR083039).

